# Chromosomal variability in a clonal crop: Somaclonal change follows the emergence of triploid saffron crocus

**DOI:** 10.64898/2026.05.04.722608

**Authors:** Abdullah El-nagish, Manoj Kumar Dhar, Ludwig Mann, Ruifang An, Andreas Houben, Frank R. Blattner, Dörte Harpke, Tony Heitkam

## Abstract

**Background:** Saffron crocus (*Crocus sativus*) is the source of saffron, the most expensive spice in the world. It evolved about 3000 years ago as a sterile triploid clone in Greece. Since then, saffron has spread across the globe, where regionally distinct practices of saffron cultivation have developed. Despite differences in morpho-physiological traits, genetic variability is low, if present at all. Here, we aim to resolve chromosomal and sequence-associated variability across saffron crocus cultivars from the crop’s main cultivation areas in Africa, Asia and Europe.

**Methods:** We used genome-wide DNA polymorphisms obtained through genotyping-by-sequencing (GBS) of 33 saffron and 14 closely related *Crocus* accessions, which we place into a phylogenetic context. For karyotyping, we compare nine saffron accessions by multi-color fluorescent *in situ* hybridisation (FISH) with repetitive DNA probes.

**Key results:** Phylogenetic analyses confirmed the single origin and clonal nature of all saffron accessions. We detected slight DNA differences among saffron crocus genotypes, which were minor compared with those in wild *C. cartwrightianus* populations. Still, the Iranian saffron accessions form a genetically very narrow group that differs from the other proveniences in population genetic analyses. However, chromosomes of some saffron accessions display variable FISH signals, likely resulting from gains and losses of tandemly repeated DNA.

**Main conclusions:** Based on the high genetic identity and small karyotypic differences, we confirm the clonal origin of the saffron accessions. Nevertheless, as we detected small and regional chromosomal variability, we conclude that at least four somaclonal saffron lineages emerged after saffron’s origin.

**Societal Impact Statement:** For millennia, many cultures developed cultivation practices and regional crop varieties. A notable case is saffron, the world’s most expensive spice that is harvested from stigmas of saffron crocus. This flower crop arose 3000 years ago in a singular genome triplication event and since then spread clonally across the globe. By identifying genetic and chromosomal variability in clonal saffron accessions, we highlight regional diversity, support the preservation of traditional knowledge, and underscore the risk of relying on only one clonal lineage. This informs strategies for saffron cultivation, linking cultural heritage with modern genomics to address biodiversity, evolution, and food security.

## Introduction

*Crocus sativus* L., known as ‘saffron crocus’ or ‘saffron’, is an economically important crop plant with culinary, medical, and cosmetic uses. Its stigmas are manually harvested from flowers in a labor-intensive process and dried to produce saffron, the world’s most expensive spice (Colapietro et al., 2019; Cerdá-Bernad et al., 2022). The major saffron-producing region is Iran, accounting for about 90% of the annual global harvest. Nevertheless, saffron use has also a long tradition in many countries around the Mediterranean Sea and on the Indian subcontinent. There, the plants are mainly grown at elevations above 1500 m a.s.l., reaching up to 1800 m in the temperate regions of Kashmir. Generally, the environment and cultivation strongly affect saffron quality, with pronounced temperature differences between day and night during the harvesting season being considered as a prerequisite for high saffron quality.

The evolutionary origin of saffron crocus has long been debated (Caiola and Canini, 2010), with diverse progenitor species and areas of domestication being postulated (Nemati et al., 2018). Phylogenetic analyses indicated that *C. cartwrightianus*, a species from the Aegean islands and adjacent mainland Greece, is the closest wild relative of *C. sativus* (Nemati et al., 2018). Phylogeographic analyses of *C. sativus* and *C. cartwrightianus* populations (Nemati et al., 2019), together with comparative fluorescent *in situ* hybridisation (FISH) to infer their karyotypes (Schmidt et al., 2019), pinpointed the wild populations in the Attica area of mainland Greece as progenitors of the saffron crocus. In this area, as well as on the island of Santorini, even today *C. cartwrightianus* is harvested as ‘wild saffron’ and used as a spice by the local population (Blattner, personal observation). Over 3600 years ago, in the Bronze Age, the diploid wild species was already cultivated in Minoan saffron gardens (Negbi and Negbi, 2002; Kazemi-Shahandashti et al., 2022), which was most probably the starting point for the selection of the extremely rare triploid variant, which we know today as *Crocus sativus, “*saffron*”*. This autotriploid’s sterility allowed it to genetically retain and to ‘freeze’ the seemingly very favorable aromatic characteristics and allelic composition of this individual. Further vegetative propagation through daughter corms resulted in the spread of this single clone, and resulted in the expansion of saffron cultivation out of the Aegean archipelago (Caiola and Canini, 2010; Nemati et al., 2019; Schmidt et al., 2019; Kazemi-Shahandashti et al., 2022).

Saffron’s triploidy (2*n* = 3*x* = 24; Brandizzi and Grilli Caiola, 1998) is the main reason for its infertility. It results in a disrupted meiosis due to flawed chromosome pairing (Chichiriccò, 1984; Rashed-Mohassel, 2020). As a consequence of pollen sterility and the presence of a self-incompatibility system, sexual reproduction is unattainable and vegetative propagation by daughter corms is considered the only way for saffron to reproduce (Caiola and Canini, 2010). This prevents improvement through breeding, as vegetative propagation by corms does not generate genome variations that usually occur through recombination during sexual reproduction.

Intriguingly, despite saffron’s clonality, phenotypic differences among accessions from different geographic regions were observed, including differences in vegetative growth size, tepal appearance, stigma count, pigmentation intensity, and aroma. Whether such differences could be attributed to epi- and/or genetic factors remains unresolved (Rubio Moraga et al., 2009; Fluch et al., 2010; Siracusa et al., 2013). Molecular marker-based studies, including simple sequence repeats (SSRs), amplified fragment length polymorphisms (AFLPs), inter-retroelement amplified polymorphism (IRAP) and expressed sequence tag-derived SSRs, have identified very low levels of genetic variation in *C. sativus* (Fluch et al., 2010; Siracusa et al., 2013; Nemati et al., 2014; Alsayied et al., 2015), leading to the widely accepted notion that saffron crocus likely originated from a single triploidisation event (Rubio-Moraga et al., 2009; Fluch et al., 2010; Siracusa et al., 2013; Nemati et al., 2019, Schmidt et al. 2019; Kazemi-Shahandashti et al., 2022). However, a recent genome-wide analysis by Busconi et al. (2021) revealed a surprising abundance of single-nucleotide polymorphisms (SNPs) and high epigenetic variability among saffron accessions. These findings indicate that saffron has a higher (epi)genetic variability than previously assumed or detected. Moreover, Agayev et al. (2010) thought they had identified small chromosomal differences in a Kashmir lineage of *C. sativus,* which may indicate an accumulation of chromosomal changes over time in addition to sequence polymorphisms.

The clonal nature of the saffron crocus should, at least in theory, result in lineage-specific fixation of mutations that do not get reorganised by genetic recombination. This might allow the inference of the genealogy of extant saffron individuals and to trace them back to, if not the initial triploid, at least their most recent common ancestor. Such an approach would potentially show the spread of saffron out of Attica and would retrace the trade routes involved in saffron dispersal. With the progress in sequencing and cytogenetics (Nemati et al., 2018; Nemati et al., 2019; Schmidt et al., 2019; Busconi et al., 2021; Kazemi-Shahandashti et al., 2022), it is timely to reinvestigate the history of saffron accessions, especially those that hold culture-defining value for entire regions. As an example, saffron was cultivated in India for over 1000 years in the Kashmir and Kishtwar regions in relative geographic isolation from other saffron-producing areas (Husaini et al., 2010; Kothari et al., 2021). Also, Iran as well as Egypt look back on long traditions of saffron use, although initially wild crocus species were mainly used as dye in paintings or for textiles (see Kazemi-Shahandashti et al., 2022).

In this study, we aim to trace the somaclonal variation in saffron crocus accessions from Europe, Iran, Kashmir and Egypt. To comprehensively address clonal variation that may arise in the form of single nucleotide polymorphisms (SNPs) in unique regions, we employ genotyping-by-sequencing (GBS; Elshire et al., 2011), a reduced genomic representation approach. This will allow us to identify saffron lineages and to reconstruct their relationships. To accommodate the potentially rapidly evolving repetitive DNA regions of the saffron crop, we screened for chromosomal variability using our six-colour saffron probe set (Schmidt et al., 2019) for fluorescent *in situ* hybridisation (FISH). With this method, we can also assess chromosomal differences that occurred in the different saffron lineages. Taken together, we aim to understand the types of somatic changes that may differentiate regional saffron crocus accessions.

## Materials and methods

### Plant material

For the analysis of genomic variations through GBS, we included 45 individuals (Supplementary Table S1), consisting of 31 individuals of saffron (*C. sativus*) representing the main growing areas of the crop, 12 individuals of its diploid progenitor species *C. cartwrightianus* from the Greek regions of Attica, Crete, Kea, Naxos, and Tinos, covering the distribution range of this species in mainland Greece and the Aegean archipelago (Nemati et al., 2019), and two individuals of *C. oreocreticus*, the sister species of *C. cartwrightianus* and *C. sativus* (Nemati et al., 2018).

For FISH karyotyping of the saffron crocus, we included *C. sativus* accessions from Egypt, France, Germany, India, Iran, Spain, and The Netherlands (Table 1). As a reference karyotype, we used the Spanish accession established before (Schmidt et al., 2019).

**Table 1.** *C. sativus* accessions used for cytogenetic investigation.

| # | Species | Accession number | Origin/Supplier |
| --- | --- | --- | --- |
| 1 | *Crocus sativus* | BCU001584 | Bank of Plant Germplasm, Cuenca, Spain |
| 2 | *Crocus sativus* | N/A | Bernd Schober Company, Augsburg, Germany |
| 3 | *Crocus sativus* | AY-EG01 | Sohag, Egypt |
| 4 | *Crocus sativus* | NL-BIO-01 | Bloembollenbedrijf J.C. Koot, Egmond Binnen, The Netherlands |
| 5 | *Crocus sativus* | 160 (ex. hort GAT) | Farmer's market, Les Vans, Dept. Ardèche, France |
| 6 | *Crocus sativus* | KSH SP10-1 | Kashmir, India |
| 7 | *Crocus sativus* | KSH SP10-2 | Kashmir, India |
| 8 | *Crocus sativus* | KSH SP11-1 | Kashmir, India |
| 9 | *Crocus sativus* | Iran_3 | Mashad, Iran |

### Genotyping-by-sequencing (GBS)

GBS analyses were conducted for 45 individuals to obtain genome-wide SNPs. Twenty-two individuals were sequenced in the framework of this project. For the library preparation, 200 ng of genomic DNA was used and cut with the two restriction enzymes *Pst*I-HF (NEB) and *Msp*I (NEB). Library preparation, individual barcoding, and 110 bp single-end sequencing on the Illumina NovaSeq were performed following Wendler et al. (2014) and Ala et al. (2025). Barcoded reads from the samples were de-multiplexed using the CASAVA pipeline 1.8 (Illumina). We added raw data from 23 samples from our previous analysis (Nemati et al., 2019) and processed them alongside the newly obtained samples in all downstream steps. Adapter trimming of GBS sequence reads was performed with CUTADAPT (Martin, 2011) within IPYRAD v.0.9.87 (Eaton and Overcast, 2020), and reads shorter than 65 bp after adapter removal were discarded. GBS reads were clustered using the IPYRAD pipeline with a clustering threshold of 0.9 and at least 30 samples sharing a locus. The default settings of parameter files generated by IPYRAD were used for the other parameters except for the maximum number of alleles, where three alleles per site were allowed in a locus to account for the triploidy of *C. sativus*.

Since IPYRAD is filtering out loci with a high proportion of shared heterozygous sites, we also used STACKS v2.55 (Rochette et al., 2019) to generate an input file to analyse shared heterozygosity aimed to infer the clonality within our data set. A locus needed to be present in 70% percent of the individuals of a species to be processed.

### Phylogenetic analyses

Maximum parsimony (MP) and SVDQuarted (SVDQ) analyses of the GBS alignment data were conducted in PAUP* 4.0a169 (Swofford, 2002). For MP a two-step heuristic search was conducted as described in Blattner (2004) with tree-bisection-reconnection (TBR) branch swapping, steepest descent not in effect, and initial 1000 random addition sequences (RAS) restricting the search to 20 trees per replicate. The resulting trees were afterwards used as starting trees in a search with maxtree set to 10,000. To test clade support, bootstrap analysis was run with re-sampling 500 times with the same settings as before, except that we did not use the initial RAS step. Two analyses were conducted. One including saffron plus *C. cartwrightianus* and *C. oreocreticus*. In this analysis, *C. oreocreticus* was defined as outgroup. A second analysis used only the 31 saffron individuals to infer genetic structure within the clonal crop. Here, no outgroup was specified resulting essentially in unrooted trees. The second dataset was analysed in addition to MP also with SVDQ, evaluating all quartets, and trees were selected using Quartet Fiduccia and Mattheyses (QFM) assembly and the multi-species coalescent (MSC) as the tree model. For bootstrap support values, we used 500 re-samples with the same settings as before but evaluating 20,000 quartets only (65% of all quartets).

To infer genetic diversity in and outside saffron, pairwise genetic distances among all individuals were calculated in PAUP* using the general time-reversible model of sequence evolution (GTR) with rates following a gamma distribution with a shape parameter of 0.5.

### Analyses of population structure and heterozygosity

To test for the presence of genetic sub-structuring within *C. sativus* an assignment analysis was conducted using the model-based Bayesian clustering in the R package ‘LEA’ (Frichot and François, 2015) together with Principal Component Analysis (PCA) to identify sub-structuring of the populations. The vcf file generated by IPYRAD v.0.9.87 was filtered using VCFTOOLS v 0.1.16 (Danecek et al. 2011) removing all indels, keeping only sites with a minor allele frequency of 0.02, a minimum sequencing depth of 10 and a maximum depth of 300, SNPs present in 90% of the individuals and keeping only *C. sativus* samples. Population assignment was performed for K = 1–10 with 30 repetitions each. The K with the lowest cross entropy was considered as optimal. PCA results obtained from LEA were visualised with ‘ggplot2’ in R (Wickham, 2016).

The shared heterozygosity (SH) index was calculated to infer the degree of clonal reproduction within *C. sativus* and *C. cartwrightianus* as described by Yu et al. (2022). It is based on the proportion of shared identically heterozygous SNPs in a pair of samples. The SH index was inferred for all samples within one species and plotted with ‘ggplot2’ in R. The STACKS2 vcf file was used as input.

### Chromosome preparation, probe labeling, fluorescent *in situ* hybridisation and microscopy

We have followed our protocol for *Crocus* chromosome preparation (El-nagish et al., 2025), with the following specific parameters: Primary roots were harvested from individual corms. After synchronisation in 2 mM hydroxyquinoline for 5 h, roots were fixed in methanol:acetic acid (3:1 v/v). Roots were then digested for 2.30 h at 37 °C in an enzyme mixture consisting of 2% (w/v) cellulase from *Aspergillus niger* (Sigma C1184), 4% (w/v) 0.5% (w/v) pectolyase from *Aspergillus japonicus* (Sigma) P-3026, cellulase Onozuka R10 (Sigma 16,419), 1% (w/v) cytohelicase from *Helix pomatia* (Sigma) C-8274, 1% hemicellulase from *Aspergillus niger* (Sigma H2125), and 20% (v/v) pectinase from *Aspergillus niger* (Sigma P4716) in citrate buffer (4 mM citric acid and 6 mM sodium citrate). Single root tips were then transferred onto slides, macerated with a needle in enzyme buffer, rinsed in 45% glacial acetic acid. Before the slide dried, chromosomes were spread on a hot surface (55 °C) and fixed with methanol:acetic acid (3:1 v/v).

Satellite DNA probes (CroSat1-4; Schmidt et al., 2019) were labeled by PCR with fluorescently labelled UTP, DY415-dUTP, 495-dUTP, 547-dUTP, and 647-dUTP (Dyomics GmbH, Jena, Germany). Sugar beet ribosomal gene probes pXV1 harboring the 5S rRNA gene, including spacers (Schmidt et al., 1994; Paesold et al., 2012) and 18S-2 containing a part of the 18S-5.8S-25S rRNA gene (Sielemann et al., 2022) were also labeled by PCR with digoxygenin-dUTP detected by antidigoxigenin–fluorescein isothiocyanate (FITC; both from Roche Diagnostics), biotin-11-dUTP (Dyomics GmbH, Jena, Germany) detected by Streptavidin-Cy3 (Sigma-Aldrich) respectively.

Prior to FISH, according to the amount of cytoplasm visible under a light microscope, slides were treated with 100 μg/ml RNase in 2× SSC for 20 min, followed by 200 μl of 10 μg/ml pepsin in 10 mM HCl for 30 min. The hybridisation and rehybridisation procedures were performed as described previously (Schmidt et al., 1994). The chromosome preparations were counterstained with DAPI (Honeywell, Charlotte, NC, USA).

Slides were examined with a Zeiss Axioimager M1 UV epifluorescence microscope with appropriate filters, and equipped with an ASI BV300-20A camera coupled with the APPLIED SPECTRAL IMAGING software (Applied Spectral Imaging, Carlsbad, CA, USA). Finally, the images were processed with ADOBE PHOTOSHOP CS5 (Adobe Systems, San Jose, CA, USA) using only contrast optimisation, Gaussian and channel overlay functions affecting the whole image equally.

## Results

To determine whether there is somaclonal variation among saffron crocus lines, we used Kashmir saffron as a starting point. From this region in India, we collected four accessions to analyse potential variations already among the genotypes from a single area. To provide further reference points, we complemented our analyses with several additional saffron accessions from regions around the globe. We combined two approaches, one focusing on genomic change in the form of single nucleotide polymorphisms (SNPs) and a second one targeting chromosomal variability in the form of tandem repeat localisation and repeat array expansion. First, on a larger scale, we used the high-throughput approach of genotyping by sequencing (GBS) to analyse the genomic variation of all 33 saffron accessions. This approach focuses on variation within unique genomic regions. Second, to also take into account the repeat fraction and the chromosomal arrangements within the saffron accessions, we complemented these efforts with repeat-based FISH karyotyping for representative saffron crocus samples from five locations.

### The GBS analysis supports the single origin of saffron crocus, highlighting its narrow genetic base

To understand the genetic variation within *C. sativus*, we performed GBS and analysed the SNP data with phylogenetic and population genetic methods. We included the diploid progenitor species *C. cartwrightianus* with at least two individuals of five different populations to be able to compare the genetic diversity within *C. cartwrigthianus* and *C. sativus* and show the phylogenetic affiliation of saffron. *Crocus oreocreticu*s, the closest relative of *C. cartwrigthianus* and *C. sativus*, served as an outgroup. In total, our GBS-derived data matrix consisted of 8,946 loci with 25.3% missing data. The concatenated characters resulted in an alignment of 999,883 bp where 98% of the positions were constant. Out of the 19,790 variable characters, 9,289 were parsimony-informative. The MP analysis resulted in 12 equally parsimonious trees of 29,767 steps in length, with a consistency index (CI) of 0.67 and a retention index (RI) of 0.64. Differences among the 12 MP trees were restricted to the relative positions of the individuals within the monophyletic group of Iranian *C. sativus*.

The phylogenetic analysis placed all saffron accessions within *C. cartwrightianus*, as sister of the group of *C. cartwrightianus* individuals from Attic populations. Bootstrap support values along the backbone of the tree are high, and saffron as well as the clade of individuals from Attica, together with saffron, obtained 100% support, each. Within saffron, genetic differences are very low in comparison to the diversity present in and among *C. cartwrightianus* populations, as indicated by branch-length differences in the tree (Fig. 1). Thus, the genetic distances between the two closest individuals from the different *C. cartwrightianus* populations in the analysis are already two orders of magnitude larger (0.00317 between two individuals from Crete) than the largest distance within all *C. sativus* individuals (0.00009 between the Dutch individuals). This is consistant with the clonal nature of worldwide cultivated saffron individuals, all going back to a single origin.

**Figure 1.**
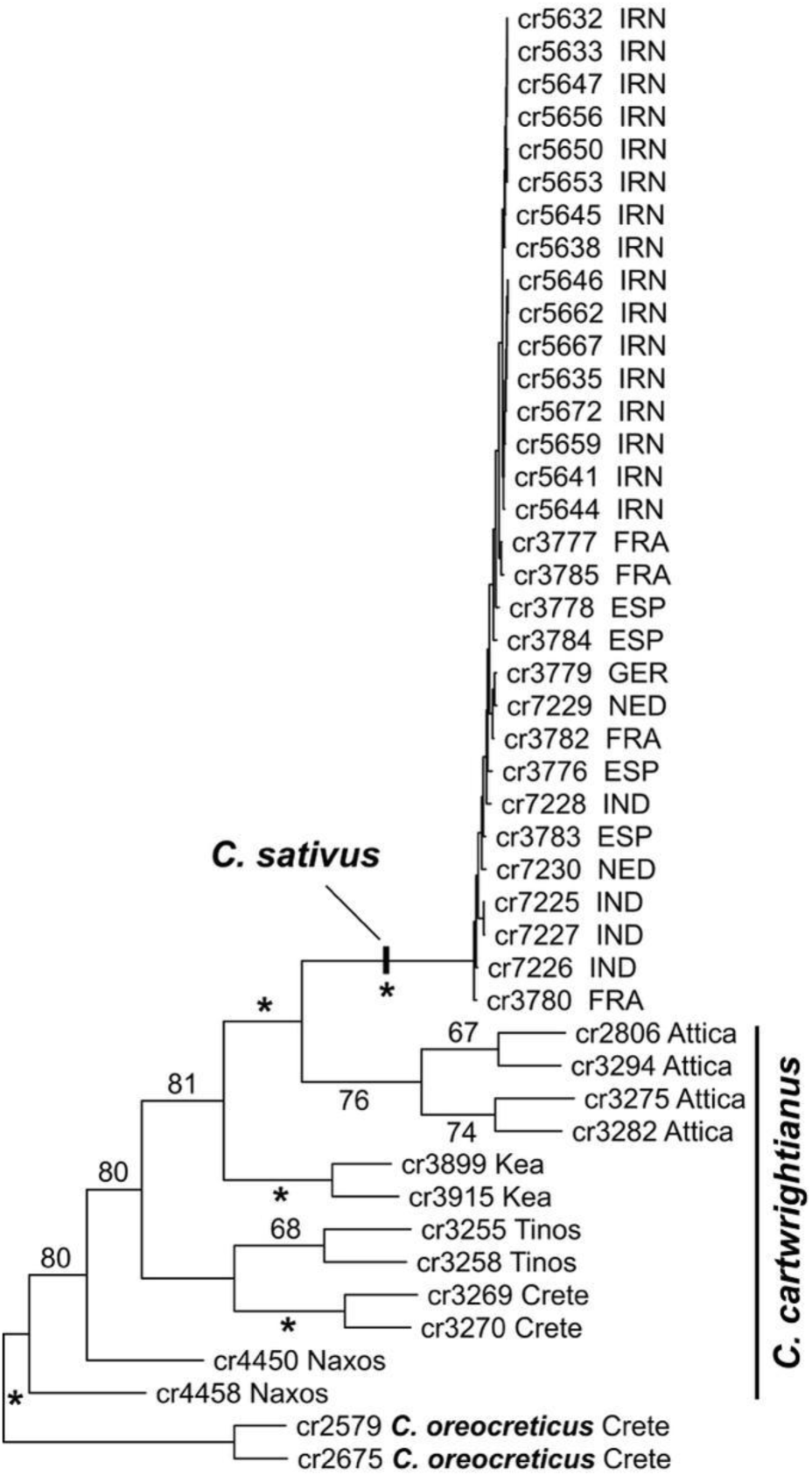
Phylogenetic analysis of *Crocus* GBS data detects only minor differences between saffron accessions. Shown is one out of 12 equally parsimonious trees of an MP analysis. Differences among these trees concern only the relative positions of some of the Iranian saffron individuals. *Crocus oreocreticus* was defined as an outgroup, numbers along the branches depict bootstrap support values (%) with 100% support indicated by asterisks. For saffron the countries of origin are given for all analysed individuals, while in *C. cartwrightianus* the respective Greek regions are provided. Low genetic differences among saffron cultivars, as indicated by very short branch lengths, are in accord with the single origin and clonal nature of the crop.

To search for genetic structure within saffron, we conducted GBS-based analyses using phylogenetic and population genetic approaches. The dataset for the 31 saffron individuals had an alignment length of 1,689,948 bp with 99.8% of constant positions. Of the 2893 variable characters 716 were parsimony informative. The MP analysis resulted in 38 equally parsimonious phylogenetic trees of 3622 steps length (CI = 0.80, RI = 0.55). Differences among the trees were restricted to the relative positions of the individuals within the monophyletic and genetically narrow group of Iranian saffron. The strict consensus of these trees (not shown) is compatible with the SVDQ tree (Fig. 2A). The Iranian individuals form a strongly supported clade in both analyses. In contrast, the individuals from Europe and Asia did not form clades according to their country of origin but occur in mixed groups. However, their corms were obtained in part in single batches from the same growers (e.g., French or Dutch materials). Even the Kashmiri accessions are not monophyletic although they are thought to have been cultivated in relative isolation for a long time period.

**Figure 2.**
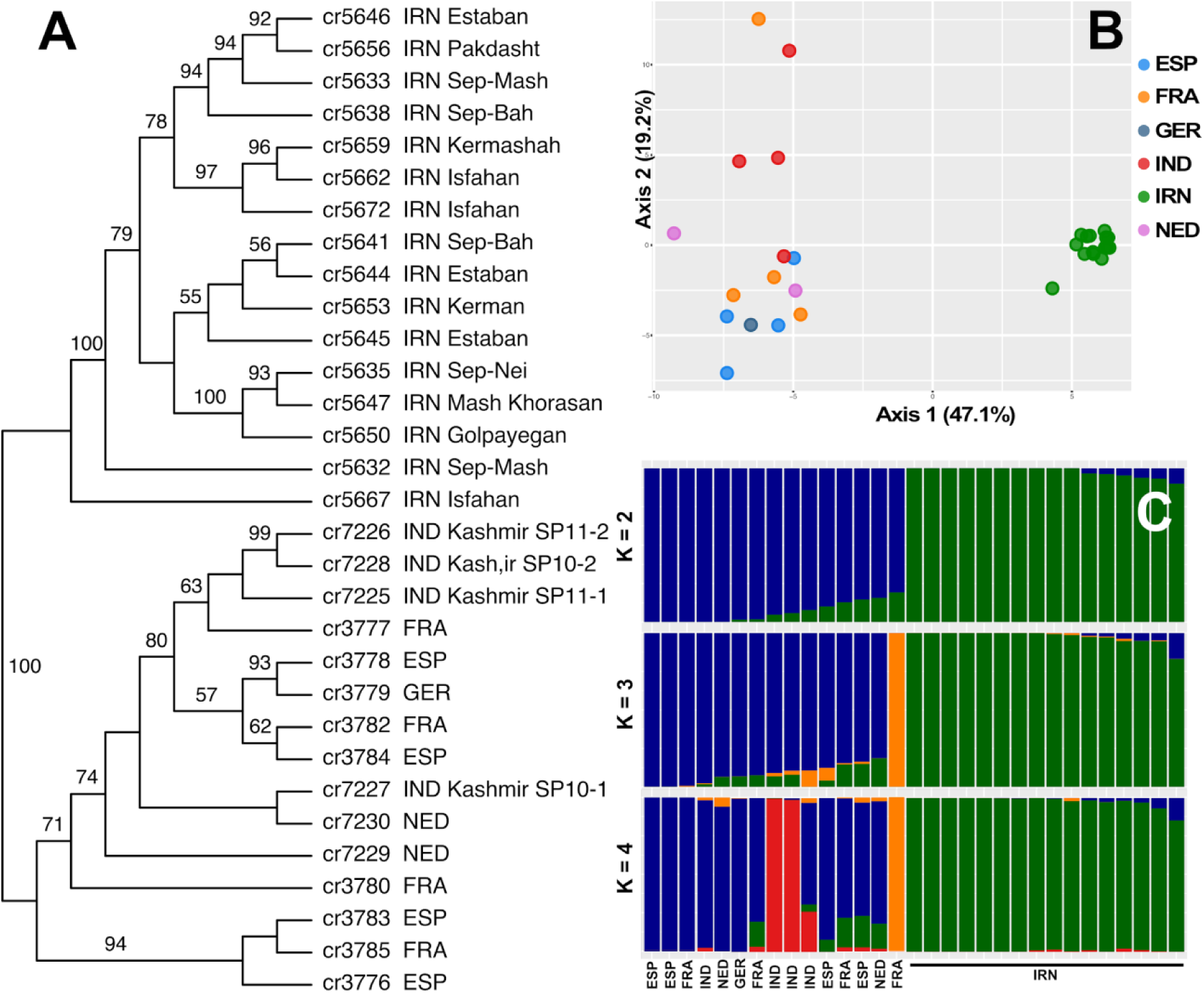
GBS-based phylogeographic and population genetic analyses of saffron (*C. sativus*) individuals found weak genetic substructuring within the clonal species. (A) Unrooted SVDQ phylogenetic tree of 31 saffron individuals from Europe, Near East, and Asia. The countries of origin of the materials are given and for the Iranian individuals also cultivation regions. Numbers along branches provide bootstrap values >50%. Have in mind that branch lengths do not reflect genetic diversity in this cladogram. **(B)** PCA plot of the first two axes explaining 66.3% of the detected genetic variation. PCA and phylogenetic analysis indicate that the Iranian saffron cultivars are a genetically rather narrow group within the species, while genetic diversity, although overall rather limited due to the clonal nature of the species, is larger in the European and Kashmiri stock of accessions. **(C)** Population assignment analyses obtained with LEA for K = 2 to K = 4.

The results of the phylogenetic analysis are closely reflected by PCA and the population structure analysis (Fig. 2). PCA separates Iranian saffron from the other accessions along the first axis and indicates them as genetically uniform. Within the non-Iranian individuals, no clear geographic structure is visible although they are separated along PCA axis 2 (Fig. 2B). In population assignment analyses the Iranian accessions are distinct from all others at K = 2. At K = 3 an individual from France is separated, while at K = 4 a split within the Kashmiri saffron was detected (Fig. 2C), reflecting the result of the SVDQ analysis and showing again that Kashmiri material is genetically not uniform.

The pairwise shared heterozygosity within saffron was found to be above 0.82 (0.82–0.96) in all possible pairing combinations, supporting that they are clonally propagated, sharing the same origin. In contrast, *C. cartwrightianus* samples shared only up to 0.28 (0.19–0.28) heterozygous sites and *C. oreocreticus* shared 0.30 heterozygous sites (Figure S1).

### Saffron lineages can be differentiated on the chromosomal level: Orthologous chromosomes of saffron crocus diverged in their repetitive DNA composition

To study the chromosomal variability between saffron accessions from different geographic regions, we performed comparative physical mapping of six repetitive DNAs by multi-color FISH. The probes include the four satellite DNAs (satDNAs) CroSat1 to CroSat4 as well as the 5S and 18S-5.8S-25S rRNA genes (Figures 3, S2-S4; Table 2). This probe set enables discrimination of all saffron chromosomes and resolves the chromosomes within triplets (Schmidt et al., 2019).

**Figure 3:**
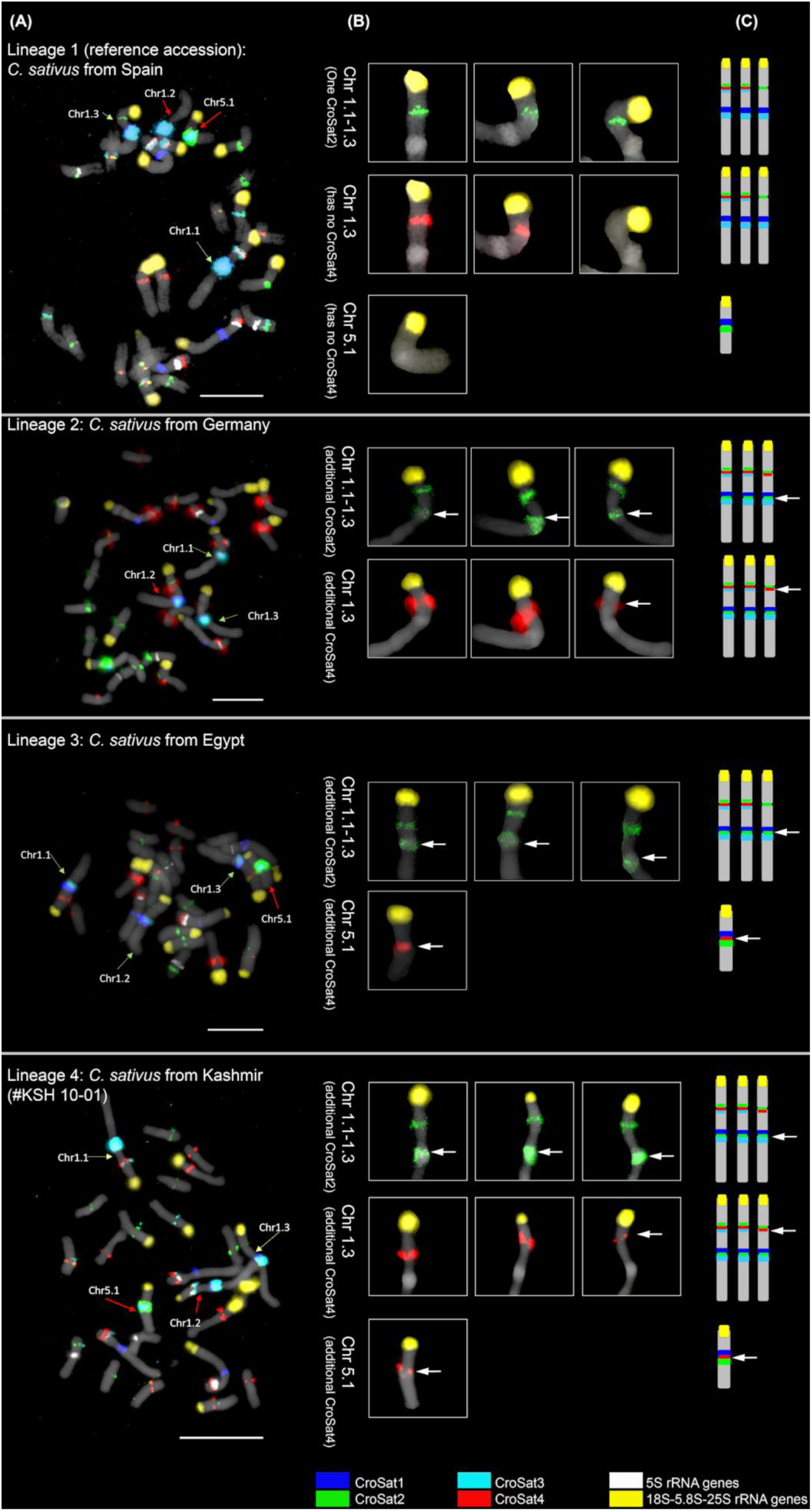
Fluorescent *in situ* hybridisation (FISH) of mitotic chromosomes of four different *C. sativus* lineages from Spain, Egypt, Germany, and Kashmir (lineage numbers are shown on the panel). DAPI-stained chromosomes are shown in gray. FISH of different *C. sativus* accessions (A); Close up of chromosomes 1.1-1.3 and chromosome 5.1 (SatDNA names given in each panel) showing the number of signals for each satDNA (B); FISH karyotype of *C. sativus* (C). Probes used are CroSat1 (blue), CroSat2 (green), CroSat3 (aqua), CroSat4 (red), 5S rRNA genes (white) and 18S-5.8S-25S rRNA genes (yellow). Arrows indicate extra signals found compared to the reference accession. Bar is 10 µm.

**Table 2.**
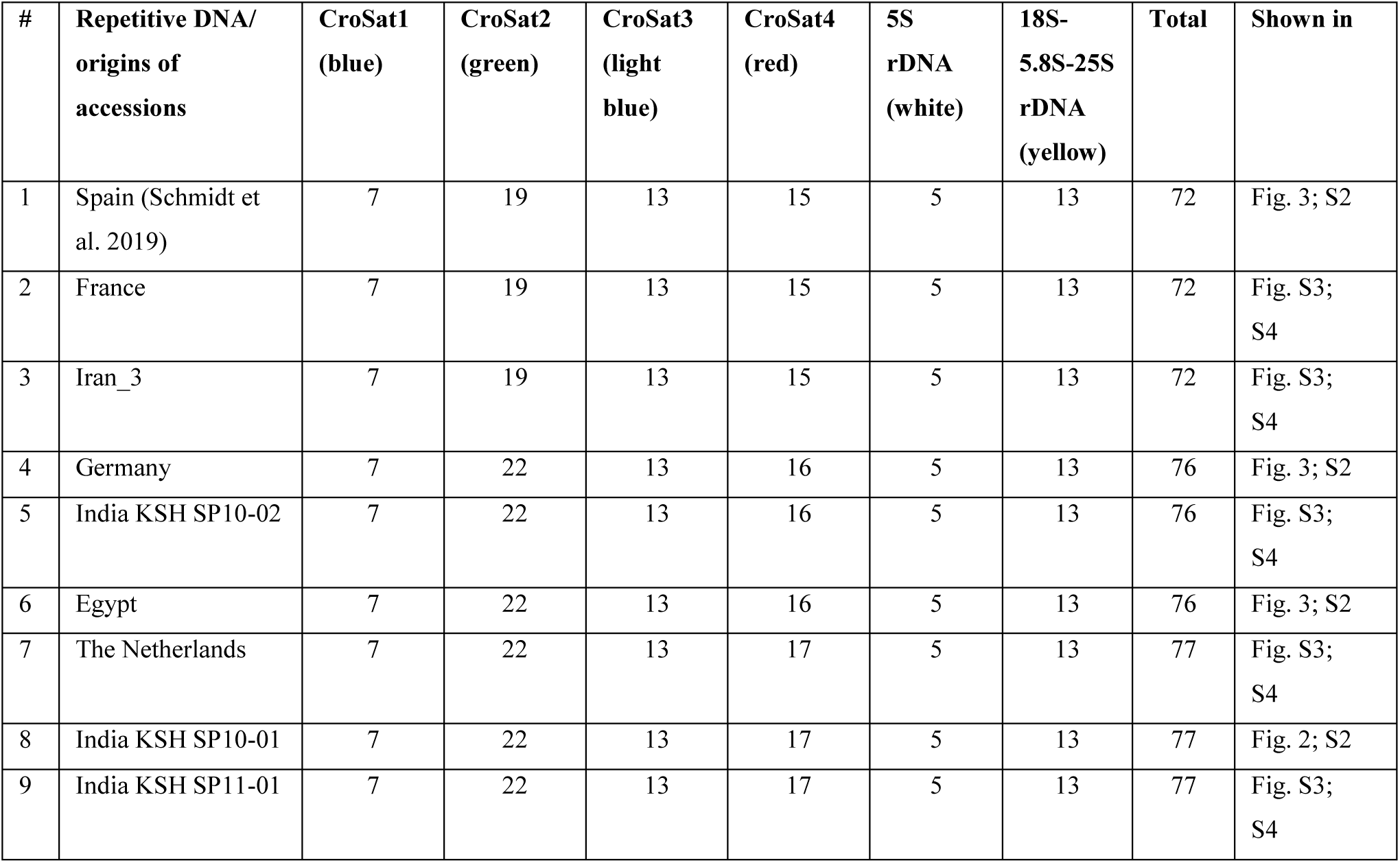
Hybridisation patterns of cytogenetic landmarks in different accessions of *C. sativus*.

| # | Repetitive DNA/<br>origins of<br>accessions | CroSat1<br>(blue) | CroSat2<br>(green) | CroSat3<br>(light<br>blue) | CroSat4<br>(red) | 5S<br>rDNA<br>(white) | 18S-5.8S-25S<br>rDNA<br>(yellow) | Total | Shown in |
| --- | --- | --- | --- | --- | --- | --- | --- | --- | --- |
| 1 | Spain (Schmidt et al. 2019) | 7 | 19 | 13 | 15 | 5 | 13 | 72 | Fig. 3; S2 |
| 2 | France | 7 | 19 | 13 | 15 | 5 | 13 | 72 | Fig. S3;<br>S4 |
| 3 | Iran_3 | 7 | 19 | 13 | 15 | 5 | 13 | 72 | Fig. S3;<br>S4 |
| 4 | Germany | 7 | 22 | 13 | 16 | 5 | 13 | 76 | Fig. 3; S2 |
| 5 | India KSH SP10-02 | 7 | 22 | 13 | 16 | 5 | 13 | 76 | Fig. S3;<br>S4 |
| 6 | Egypt | 7 | 22 | 13 | 16 | 5 | 13 | 76 | Fig. 3; S2 |
| 7 | The Netherlands | 7 | 22 | 13 | 17 | 5 | 13 | 77 | Fig. S3;<br>S4 |
| 8 | India KSH SP10-01 | 7 | 22 | 13 | 17 | 5 | 13 | 77 | Fig. 2; S2 |
| 9 | India KSH SP11-01 | 7 | 22 | 13 | 17 | 5 | 13 | 77 | Fig. S3;<br>S4 |

The baseline for our comparison is the saffron reference accession from Spain for which we have established the saffron reference karyotype (Schmidt et al., 2019; Figures 3, S2, first row). For this Spanish accession, the total number of signals is 72, detailed as following:

- 7 hybridisation sites on 7 individual chromosomes for CroSat1;
- 19 sites on 19 individual chromosomes for CroSat2, all in (peri)centromeric location;
- 13 sites on 11 chromosomes for CroSat3, with chromosomes 1.1 and 1.2 having two repeat arrays each;
- 15 sites on 15 chromosomes for CroSat4, all in (peri)centromeric location;
- 5 hybridisation sites on 5 individual chromosomes for the 5S rDNA; and
- 13 hybridisation sites on 13 individual chromosomes for the 18S-5.8S-25S rDNA.

To understand if the other selected saffron accessions differ in their FISH pattern, we comparatively mapped the six probes along their chromosomes. In total, we investigated individuals from the three Kashmir accessions as well as individuals from five geographically distant regions, obtained from Germany, the Netherlands, France, Iran and Egypt. These were then compared to the published saffron FISH karyotype (Schmidt et al. 2019; Figures 3, S2, first row). Overall, we counted 72 to 77 FISH signals, depending on the saffron accession (Table 2). Small signal number variations were found between different saffron genotypes. Nevertheless, most of the signals were shared among all accessions, indicating their origin from a single clonal line.

Whereas hybridisation of four probes (CroSat1, CroSat3, both rDNAs) yielded uniform signal patterns as published (Schmidt et al. 2019), variation occurred in CroSat2 and CroSat4, both in landmark number and position along the chromosomes of the first triplet and chromosome 5.1 (Figure 3).

In contrast to the 19 signals counted after CroSat2 hybridisation to the reference accession, most other saffron accessions displayed three additional CroSat2 signals (Figures 3; S2-S4; green signals). Interestingly, these were not near the centromere, but located intercalarily in the brightly DAPI-stained knob region along the long arms of chromosome triplet 1 (Figures 3, S3). Only the reference accessions from Spain (Schmidt et al. 2019), France and Iran corresponded to the published reference karyotype; the others showed the additional CroSat2 signals.

For CroSat4, the number of signals ranged from 15 to 17 among saffron accessions. The three accessions from Kashmir, the German and the Dutch accessions showed a CroSat4 array on chromosome 1.3 that is not found in the other accessions. An additional CroSat4 polymorphism occurred in the knob region of the heteromorphic chromosome 5.1: Accessions from Egypt, the Netherlands and two from Kashmir (KSH SP10-1, KSH SP11-1) showed an array of CroSat4 that is not found in the other accessions.

Comparing the pattern and number of hybridisation sites of each individual accession to the reference (Figures 3, S2; row 1; Schmidt et al., 2019) revealed more about the chromosomal variability in saffron accessions:

- The accessions from France and Iran showed the same pattern and number of hybridisation sites as the reference karyotype from Spain.
- The Egyptian accession exhibited a pattern that was not seen in any other accession. Compared to the reference, it exhibited similar results after hybridisation with CroSat1, CroSat3, 5S rRNA, and 18S-5.8S-25S rRNA. However, it showed additional intercalary signals for CroSat2 on all three chromosomes of the first triplet and a pericentromeric signal for CroSat4 on chromosome 5.1.
- One of the Kashmir accessions (KSH SP10-2) is comparable to the one obtained from a German source. Compared to the reference accession, both showed consistent signal patterns for CroSat1, CroSat3, the 5S and the 18S-5.8S-25S rDNA. However, both diverge from the reference accession after hybridising with CroSat2 and CroSat4. CroSat2 yields additional intercalary signals on all three chromosomes of the first triplet, whereas CroSat4 produces an additional pericentromeric signal on chromosome 1.3.
- Two of the Kashmir accessions (KSH SP10-1 and India KSH SP11-1) and one from the Netherlands have very similar signal patterns, with the total number of signals being 77. These accessions have consistent signal patterns for CroSat1, CroSat3, 5S rRNA, and 18S-5.8S-25S rRNA, but show additional intercalary signals for CroSat2 on all three chromosomes of the first triplet, a pericentromeric signal for CroSat4 on chromosome 1.3 and a pericentromeric signal for CroSat4 on chromosome 5.1.

Taken together, most cytogenetic landmarks were similar across all tested accessions. Noteworthily, all 72 published landmarks (Schmidt et al. 2019) were identified in all saffron accessions. Nevertheless, three different kinds of chromosomal variability were identified: I) chromosome triplet 1.1-1.3 may show additional CroSat2 signals; II) chromosome 1.3 may show an additional CroSat4 signal; and III) chromosome 5.1 may show an additional CroSat4 signal. Based on these differences, we can differentiate four different saffron lineages: Lineage 1 (with the three accessions from Spain, France and Iran); Lineage 2 (with the German and Indian KSH SP10-2 accessions); Lineage 3 (with the Egyptian accession); Lineage 4 (with the Dutch and Indian KSH SP10-1/KSH SP11-1 accessions) (Figure 4).

**Figure 4.**
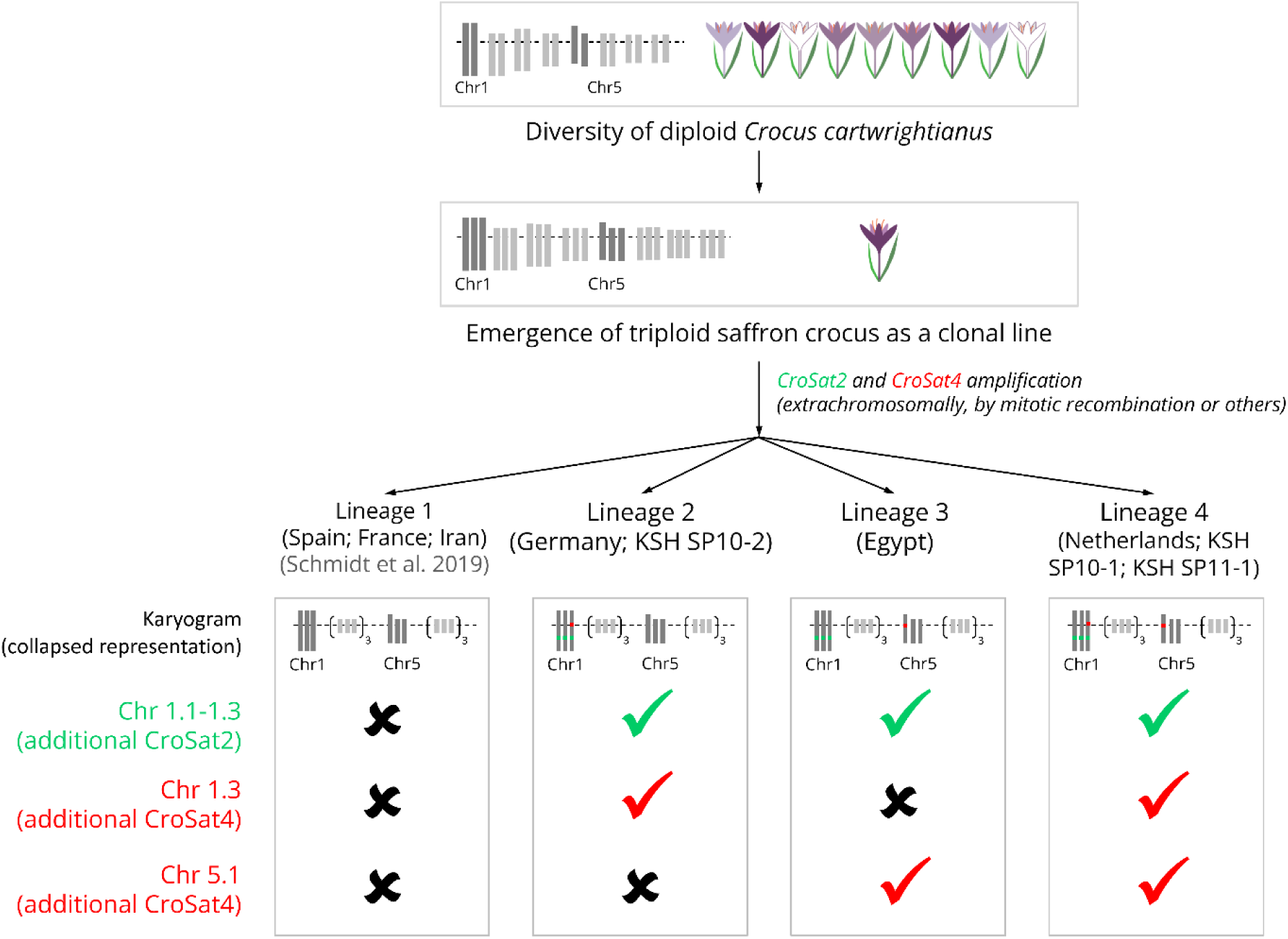
Evolutionary scenario illustrating the emergence of triploid saffron (*Crocus sativus*) as a clonal line and the subsequent accumulation of somatic chromosome mutations, leading to the split into at least four genetically distinct lines. These mutations may have altered the content of tandem repeats, specifically CroSat2 and CroSat4. Mechanisms that might be involved in driving these changes include extrachromosomal amplification followed by reintegration into chromosomes, chromosomal rearrangements such as mitotic recombination and translocation of repeats, segmental duplications, and retrotransposon-associated repeat amplification. These processes likely contributed to the expansion or loss of tandem repeats over time. Symbols (✖/✔) indicate the absence (✖) or presence (✔) of satellite DNA signals, for which green checkmarks represent presence of CroSat2 and red checkmarks of CroSat4.

## Discussion

### Chromosomal variability among saffron accessions reveals at least four saffron lineages, likely resulting from somaclonal variation

Saffron crocus is a sterile, triploid flower crop that has been propagated for over three millennia as a clonal line. High genetic similarity across all saffron crocus accessions tested so far (this report; Fluch et al., 2010; Alsayied et al., 2015; Nemati et al., 2018, 2019; Torricelli et al., 2019; Cardone et al., 2021) leaves no doubt that the cultivated triploid saffron crocus has emerged only once. Indeed, the site of saffron’s emergence was traced back to Ancient Greece and wild *C. cartwrightianus* as a sole parent-contributing species (Nemati, et al., 2019; Schmidt et al., 2019). Nevertheless, some saffron accessions are characterised by differing morpho-physiological traits depending on their region of cultivation (e.g., Torricelli et al., 2019; Cardone et al., 2021). Whether these different traits are based mainly in genetic or epigenetic change or a mix of both is currently still investigated (Busconi et al., 2018, 2021; Kazemi-Shahandashti et al., 2022).

Following a report observing a small chromosomal difference in saffron accessions from Kashmir (Agayev et al., 2010), we used three Kashmir accessions to address potential somaclonal changes using a combination of sequencing and cytogenetics. To generalise and place our results in context, we complemented these accessions with saffron from many other geographical regions. Subjecting 31 different *C. sativus* accessions to genotyping-by-sequencing, we confirmed their low genetic diversity and common origin. We also concluded that GBS only detects few statistically significant genetic differences among saffron crocuses from different regions across the globe. GBS targets unmethylated regions of the genome, which are primarily single-copy and gene-rich (Elshire et al., 2011). By employing methylation-sensitive enzymes such as *Msp*I, GBS avoids cutting at methylated cytosine residues. These methylated sites are typically associated with repetitive sequences and other non-coding regions, allowing the method to preferentially target functionally relevant genomic regions. The repetitive fraction of genomes, including transposable elements and tandem repeats, is generally considered prone to rapid evolution, often exhibiting polymorphisms or array expansions (Garrido-Ramos, 2017). As a result, variability in these regions, as well as structural differences among saffron accessions, remain “invisible” to GBS.

Nevertheless, there is the opportunity to assess variability in the repetitive DNAs by integrating a chromosomal perspective. For this, we have analysed a subset of saffron crocus accessions cytogenetically, using our 6-color repetitive DNA FISH probes (Schmidt et al., 2019). These probes differentiate the chromosomal triplets and the chromosomes within, hence, allowing to assess the chromosomal variability at high resolution. In total, we have analysed nine saffron crocus accessions obtained from breeders in Asia, Europe and Africa. Importantly, we find that the large majority of FISH signals are similar among all accessions, thus confirming the singular emergence of triploid saffron crocus as a clonal line. Nevertheless, among the accessions, we detected chromosomal variability in three independent instances, i.e. additional CroSat2 signals on chromosomes 1.1-1.3; an additional CroSat4 signal on chromosome 1.3; an additional CroSat4 signal on the heteromorphic chromosome 5.1.

Regarding other chromosomal regions, such as the nucleolus organiser regions (NORs) marked by the 18S-5.8S-25S rDNA probe, we observe a uniform number of signals across all accessions. We also confirm the observations of Agayev et al. (2010), who detected variability in NOR expansion; here, we saw this as variability in signal intensity and signal shape. Nevertheless, as variability in the size and position of the NOR has been reported in plants (Schubert and Wobus, 1985; reviewed in Garcia et al. 2024), we are not basing our conclusions on those. Instead, we will rely on the uncommon presence/absence of satellite DNA signals as laid out above.

Based on the combinations of the three sites of chromosomal variability, we conclude that at least four different saffron lineages evolved somaclonally after the emergence of triploid saffron crocus (see Figure 4). We suggest that *C. sativus* spread to different areas around the globe and accumulated mutations/chromosome alterations, such as amplification and elimination of satDNAs, over thousands of years of vegetative propagation, giving rise to multiple saffron crocus lineages. There is a high likelihood that testing additional saffron crocus accessions will reveal additional somaclonal variability.

Compared to other clonally propagated crops, we know only very little about the accumulation of genetic changes after the emergence of the triploid saffron crocus: All marker-based approaches yielded no statistically relevant variation (see cited sources above). The only variability detected by molecular gel-based markers, where genetic variability was estimated as 7.8% (Gautam and Bhattacharya, 2021), showing that clonal changes are feasible and detectable in saffron. Then, in 2021, the first comparative genome sequencing showed that there are SNPs between accessions, also among potential genic regions (Busconi et al., 2021). Based on these reports, SNP markers were developed that were able to identify polymorphisms between saffron accessions (Lachheb et al., 2023). Still, it is difficult to draw definitive conclusions: At the time of the cited analyses, there was no annotated saffron reference genome available, and many factors of insecurity may have disturbed the analysis, such as the high amount of repetitive DNA (Schmidt et al., 2019) that had not yet been annotated and the history of whole-genome duplications in saffron’s past (Xu et al., 2024) that had not yet been taken into account.

However, there is a range of crops that may provide further guidance towards contextualising the detected somatic changes, such as plants propagated through tissue culture, grafts or explants (e.g., bananas, oil palm, apple, potato or strawberry) or in apomictic plants (reviewed in Bairu et al., 2011; Roux et al., 2021; Sánchez-Romero, 2024). Similar to saffron, most genomic insights are based on molecular markers (e.g. Vroh-Bi et al., 2011), and information regarding the somaclonal variability is hard to grasp: For example, combining single sequence repeat markers and cytogenetics revealed that the triploid East African highland bananas emerged from a single clone (Němečková et al., 2018), but the extent of their genetic variation remained inaccessible. Now, the first sequencing reports are emerging: For clonal triploid bananas, SNPs were detected from RNA sequencing to estimate heterozygosity in the clonal lines and to develop SNP markers for the detection of somaclonal variants (Hou et al., 2022). In the crop model rice, whole genome sequencing gave insight into the molecular basis of these changes. The rice somaclonal variations appeared as large duplications and deletions, and as transpositions of the *Tos17* element and as frequent SNPs (Miyao, 2024).

Saffron’s morpho-physiological trait variation can also result from epigenetic changes (Busconi et al., 2018; Busconi et al., 2021; Kazemi-Shahandashti et al., 2022) and the effects of the surrounding microbiome (Ramandi et al., 2023). High epigenetic variability has already been determined for several saffron ecotypes, which were not unitised by cultivation in the same field (Busconi et al., 2018; Busconi et al., 2021).

Here, we detected large chromosome-scale genetic changes that separate different saffron crocus lineages from each other. Changes in chromosome number and structure have long been associated with somaclonal variation during *in vitro* plant propagation (e.g. Gupta, 1998). Reasons for this have been speculated to relate to the different progression times of hetero- and euchromatin through the somatic cell cycle (Lee et al., 1988; Bairu et al., 2011). Similar processes may also occur during the natural clonal propagation of saffron crocus, potentially leading to the observed chromosomal variability and split into clonal lineages. In this report, we observed changes in the two highly repetitive satellite DNAs CroSat2 and CroSat4, together making up nearly 1% of the genome (Schmidt et al., 2019). The additional hybridisation signals after FISH likely result from expansion of small satellite DNA arrays that fell below the detection threshold before. Satellite DNA array expansions can be a result of mitotic recombination (Jaco et al., 2008; Zafar et al., 2017). Alternatively, new satellite DNA locations may appear via chromosomal integration of extrachromosomal circular DNA (eccDNA) and subsequent sequence amplification (Navrátilová et al. 2008; Cohen et al. 2010; Mann et al. 2022). The same processes may also lead to changes in the nucleolus organiser regions (NOR) as has been observed for saffron crocus by Agayev et al. (2010) and by us (this report). These chromosomal changes are mechanistically possible, as similar satellite DNA amplifications and dispersions have been observed in plants like *Medicago* and *Chenopodium* (Rosato et al., 2012; Belyayev et al., 2020).

Summarising, we present clear evidence for the separation of the clonal saffron crop into somaclonal lineages. We also advocate for a combination of methods (sequencing and cytogenetics) to obtain a more comprehensive picture of the scope of saffron’s variability.

### Can clonal crops such as saffron crocus be traced to a certain region? – Implications, potentials and pitfalls

We present evidence for at least four somaclonal chromosomal lineages within the saffron crop. However, the changes found cannot be ordered in a genealogical manner, which would allow defining a progression of karyotype changes through time. This might be a result of either too few lineages be analysed and that we missed important variants, or such variants might no longer be present in the extant gene pool of saffron. Our GBS data show, that genetic differences are present within saffron accessions, although they are very small in comparison to the wild species (Fig. 1) and mostly don’t provide a clear geographic pattern (Fig. 2A). They main result here is that the extant Iranian cultivars are differing from all other analysed saffron lineages (Fig. 2) and, although representing the main production area of saffron, are genetically very narrow (Fig. 2B) even when compared to the other proveniences of the crop. Our phylogenetic analysis indicates that the Iranian accessions might be derived within *C. sativus* (Fig. 1) and that Kashmiri saffron is not related to the Iranian cultivars, but has an independent origin within or together with the stock of today’s European lineages.

Now, we can try to answer more general questions:

I. Is it possible to trace somaclonal variation? We here provide the first insight into the chromosomal basis of somaclonal changes in saffron crocus. Nevertheless, we cannot yet give insight into which saffron variant emerged first, and if one saffron variant resulted from another. We suggest that a larger panel of saffron accessions may enable the reconstruction of saffron’s history after the initial triploidisation event 3000 years ago.
II. Can saffron crocus variants be linked to a certain region? Circling back to the three Kashmir accessions that served as the starting point of our analysis, we can conclude that they have chromosomal characteristics that separate them from other saffron lineages. Yet, not from all, as we detected similar variation elsewhere. Similarly, the three Kashmir lines were not grouped together, but rather fell into two lineages. If these results reflect human intervention in the past, are results of global trade, or reflect parallel processes in clonal evolution and clonal selection, we cannot conclusively resolve. Nevertheless, the clonal genetic variability we identified here may serve as a baseline to ensure quality traits associated with regional saffron accessions, e.g., through geographical indications (GIs). GIs are intellectual property rights, which protect the name of a product with a specific geographical origin, in the light of specific qualities and/or reputation according to its particular origin (Cardoso et al., 2022). GI tags have many benefits, but also limitations and even political dimensions, as seen in the still ongoing Basmati rice controversy (Lightborne, 2003; Subbiah, 2004; Upreti, 2023). GIs are also a recurring issue for readers of *Plants, People, Planet*, as they were discussed in specific cases, such as for “Ceylon cinnamon” (Suriyagoda et al., 2021) and more generally with regard to policies for benefit sharing of natural resources in low and middle income countries (Blackhall-Miles et al., 2023).
III. What is the potential, what are the pitfalls? We show here that chromosomal variability is a source for the emergence of saffron lineages. This information has a lot of potential in saffron breeding, as it can help to inform clonal selection initiatives as they are already performed around the globe (e.g. Agayev et al., 2009; Douglas et al., 2014; El Caid et al., 2020). Similarly, the development of markers can help to generate new clonal saffron lineages, possibly *de novo* generated by chemically induced mutations. In the past, colchicine and ethyl methanesulfonate (EMS) treatments have already led to morphological variation, such as saffron with more threads and changed aroma (Samadi et al., 2022). As analyses into the diploid wild relatives deepen (Nemati et al., 2019; Schmidt et al., 2019), rebreeding of triploid saffron crocus may also emerge as an option. For other clonal crops, these kinds of breeding initiatives already exist, such as for enset (“false banana”), where triploids are bred from diploid wild germplasm, which are then maintained clonally (White et al., 2023).

Taken together, we here explored the chromosomal basis of clonal variation among saffron crocus and showed that there are at least four saffron lineages. We discussed the implications of clonal variability of saffron crocus in terms of variant tracing, protection and potential for breeding and selection.

## Acknowledgements

We acknowledge DFG funding awarded to TH (HE 7194/2-1) and FRB (BL 462/19-1) as part of the DFG sequencing call 2 (project 433081887), and to DH (HA 7550/4; project 465449547). A fellowship from the Egyptian Ministry of Higher Education was awarded to AE (Call 2019-2020).

## Author Contribution

MKD collected and initially tested the Kashmir samples. AE, RA, DH and FRB provided the remaining plant material. AE performed the cytogenetic analysis. DH and FRB performed genotyping-by-sequencing and analysed the data. AE, MKD, FRB, DH and TH wrote the manuscript. AE, MKD, AH, FRB and TH conceived and initialised the idea. TH coordinated the process. All authors discussed the results and contributed to the final manuscript.

## Data Availability Statement

GBS datasets are available under the ENA study ID PRJEB27327.

## Author information

- Abdullah El-nagish;
- Manoj Kumar Dhar;
- Ludwig Mann;
- Ruifang An;
- Andreas Houben;
- Frank R. Blattner;
- Dörte Harpke;
- Tony Heitkam;

