## Supplementary Files for "Chromosomal variability in a clonal crop: Somaclonal change follows the emergence of triploid saffron crocus"

***El-nagish et al.*** **Supporting Information**

Article title: Somaclonal changes after the emergence of triploid saffron crocus results in chromosomal variability of a clonal crop

Authors: Abdullah El-nagish, Manoj Kumar Dhar, Ludwig Mann, Ruifang An, Andreas Houben, Frank R. Blattner, Dörte Harpke, Tony Heitkam

The following Supporting Information is available for this article:

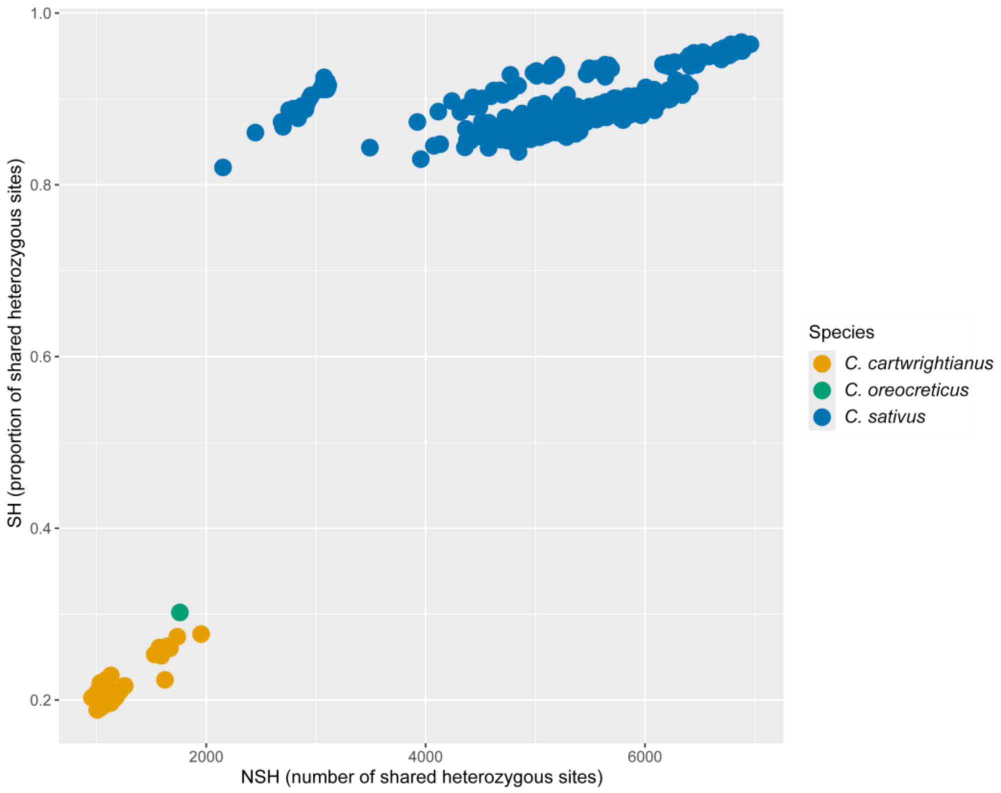


**Fig. S1:** Detecting clonemate pairs using the shared heterozygosity (SH) index for 33 *C. sativus* accessions (blue), ten *C. cartwrightianus* accessions (orange) and two *C. oreocreticus* accessions (green) based on 7250–14978 loci (average of 13777; 60856 sites). Each data point represents one pair of intra-specific samples. The SH index values of all the *C. sativus* sample pairs are close above to 0.82, meaning that all samples originated from the same clone.

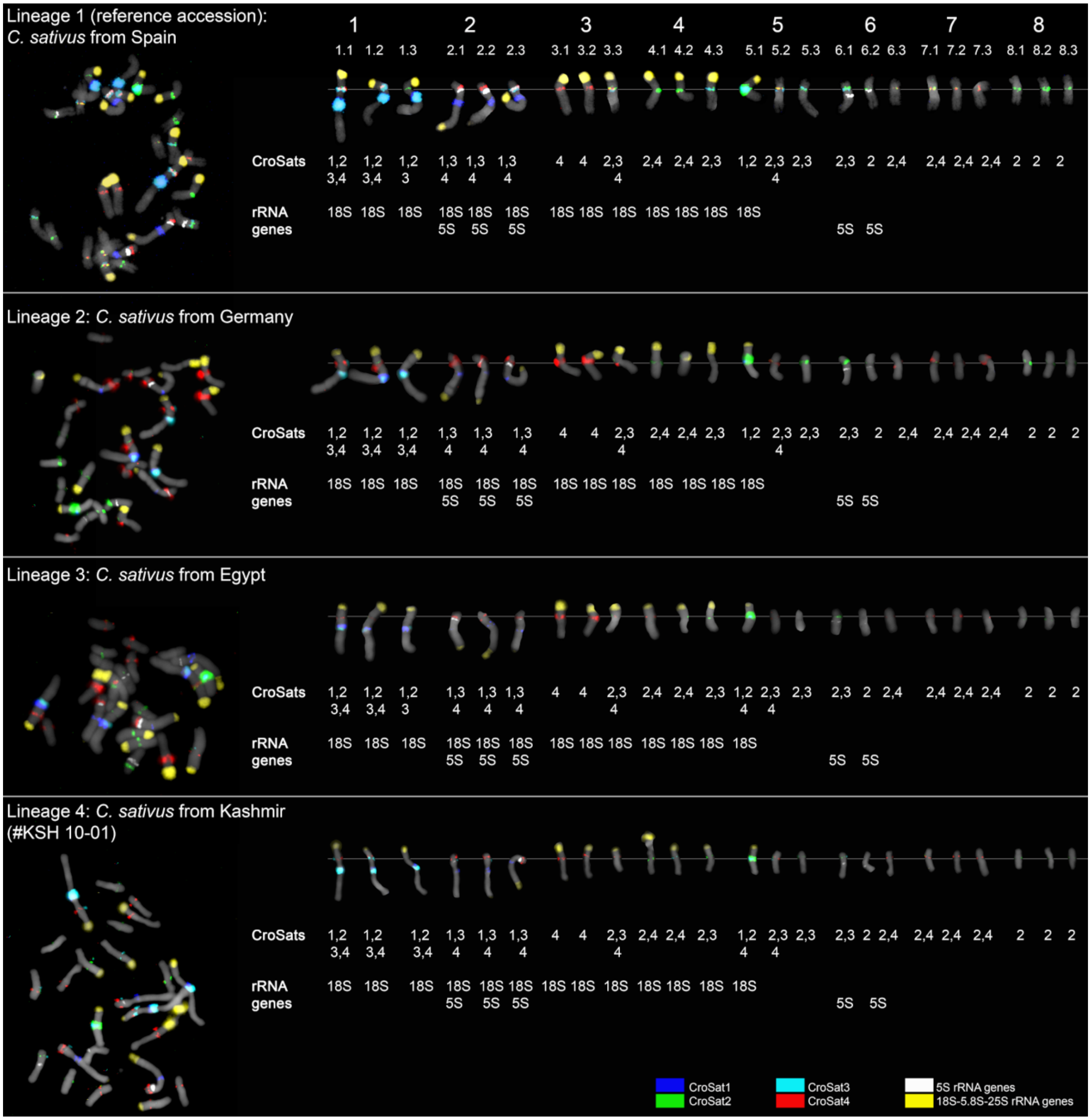


**Fig. S2:** Karyotypes of five different *C. sativus* lineages from Spain, Egypt, Germany, and Kashmir and Iran (lineage numbers are shown on the panel). DAPI-stained mitotic chromosomes are shown in gray. FISH of each *C. sativus* accession showing a mitotic metaphase (Left); and the arranged chromosome triplets (Right) showing the number of signals for each satDNA and rDNA probe carried by each chromosome. Probes used are CroSat1 (blue), CroSat2 (green), CroSat3 (aqua), CroSat4 (red), 5S rRNA genes (white) and 18S-5.8S-25S rRNA genes (yellow).

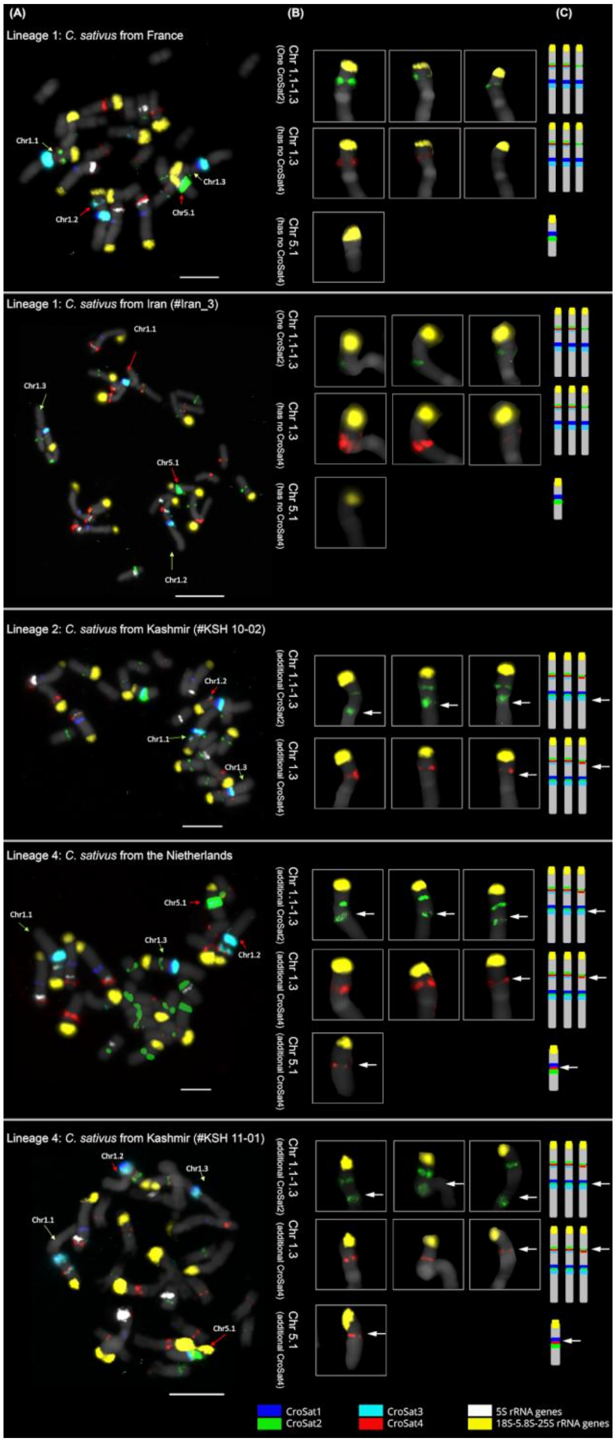


**Fig. S3:** Fluorescent *in situ* hybridisation (FISH) of the remaining different *C. sativus* lineages from France, Nietherlands, Kashmir and Iran (lineage numbers are shown on the panel). DAPI-stained mitotic chromosomes are shown in gray. FISH of a *C. sativus* accession showing a mitotic metaphase (A); Close up of Chr 1.1-1.3 and Chr 5.1 (SatDNA names given in each panel) showing the number of signals for each satDNA (B); FISH karyotype and ideogram of *C. sativus* (C). Probes used are CroSat1 (blue), CroSat2 (green), CroSat3 (aqua), CroSat4 (red), 5S rRNA genes (white) and 18S-5.8S-25S rRNA genes (yellow). Arrows indicate extra signals found compared to the reference accession. Bar is 10 µm.

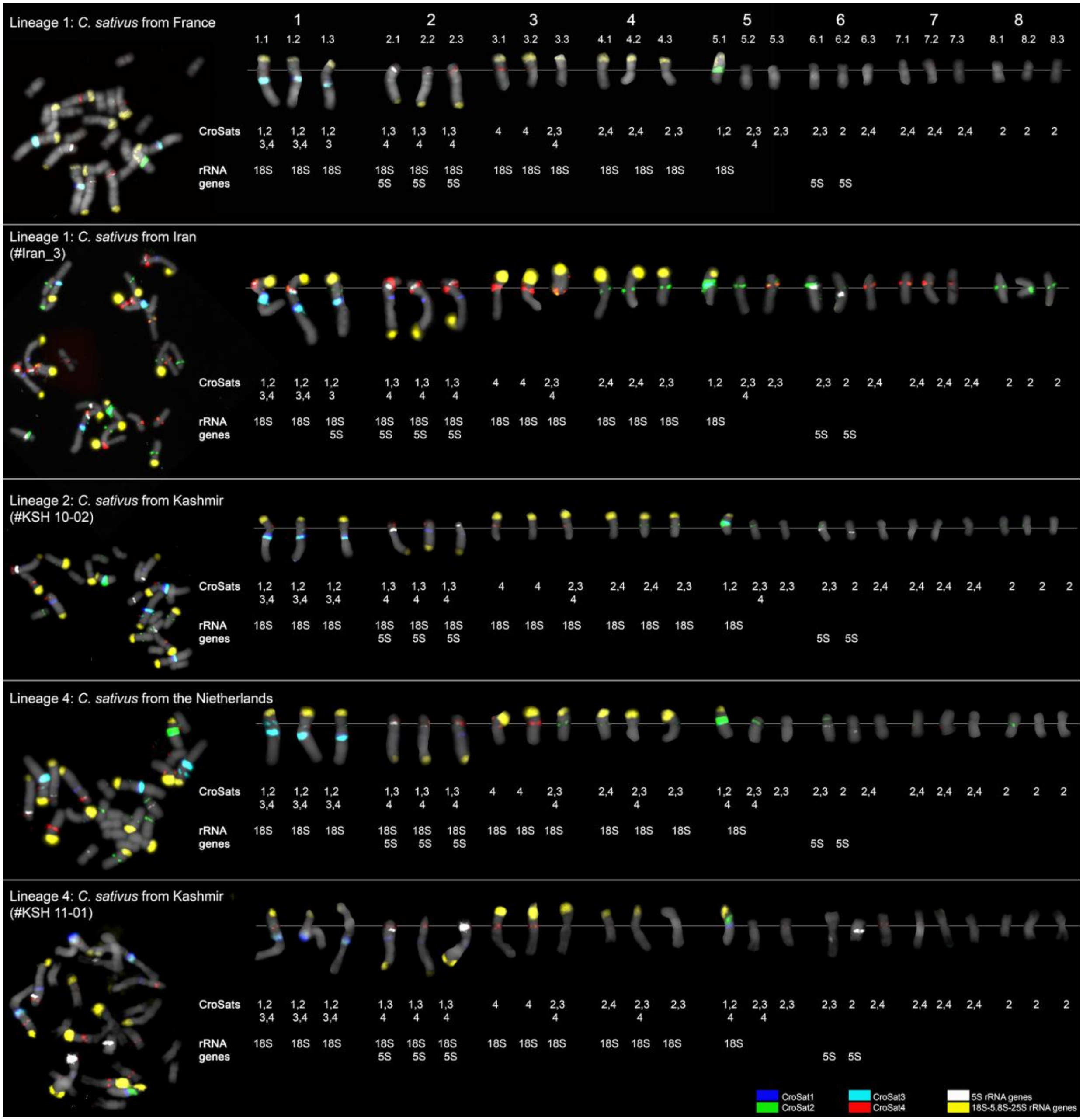


**Fig. S4:** Karyotypes for the metaphases shown in Figure S2 of the remaining *C. sativus* lineages from France, Nietherlands, Kashmir and Iran (lineage numbers are shown on the panel). DAPI-stained mitotic chromosomes are shown in gray. FISH of each *C. sativus* accession showing a mitotic metaphase (Left); and the arranged chromosome triplets (Right) showing the number of signals for each satDNA and rDNA probe carried by each chromosome. Probes used are CroSat1 (blue), CroSat2 (green), CroSat3 (aqua), CroSat4 (red), 5S rRNA genes (white) and 18S-5.8S-25S rRNA genes (yellow).

### Table S1. *Crocus* accessions used for GBS.

| Species | Accession | DNA ID | Origin |
|---|---|---|---|
| *C. sativus* L | | | |
| | Sep-Mash | cr5632 | Sepidan (origin: Mashhad), Iran |
| | Sep-Mash | cr5633 | Sepidan (origin: Mashhad), Iran |
| | Sep-Nei | cr5635 | Sepidan (origin: Neyshabur), Iran |
| | Sep-Bah | cr5638 | Sepidan (origin: Bahabad), Iran |
| | Sep-Ker | cr5641 | Sepidan (origin:Kermanshah), Iran |
| | Est | cr5644 | Estahban, Iran |
| | Est | cr5645 | Estahban, Iran |
| | Est | cr5646 | Estahban, Iran |
| | Mash_Khorasan | cr5647 | Mashhad, Iran |
| | Golpayegan | cr5650 | Golpayegan, Iran |
| | Kerman | cr5653 | Kerman, Iran |
| | Pakdasht | cr5656 | Pakdasht, Iran |
| | Kermanshah | cr5659 | Kermanshah, Iran |
| | P1_I_1 | cr5662 | Isfahan, Iran |
| | P2_I_1 | cr5667 | Isfahan, Iran |
| | P3_I_1 | cr5672 | Isfahan, Iran |
| | cp1 IND Kash | cr7225 | Kashmir, India |
| | cp2 IND Kash | cr7226 | Kashmir, India |
| | ck1 IND Kash | cr7227 | Kashmir, India |
| | ck2 IND Kash | cr7228 | Kashmir, India |
| | NL-BIO-01 | cr7229 | Bloembollenbedrijf J.C. Koot, Egmond Binnen, The Netherlands |
| | NL-BIO-01 | cr7230 | Bloembollenbedrijf J.C. Koot, Egmond Binnen, The Netherlands |

| | | | |
|---|---|---|---|
| | BCU001584 | cr3776 | Bank of Plant Germplasm, Cuenca, Spain |
| | 142-2b | cr3777 | Farmer's market, Les Vans, Dept. Ardèche, France |
| | BCU001584 | cr3778 | Bank of Plant Germplasm, Cuenca, Spain |
| | 146 | cr3779 | Saxen-Safran Stolpen, Germany |
| | 161 | cr3780 | Farmer's market, Les Vans, Dept. Ardèche, France |
| | 162 | cr3782 | Farmer's market, Les Vans, Dept. Ardèche, France |
| | BCU001584 | cr3783 | Bank of Plant Germplasm, Cuenca, Spain |
| | BCU001584 | cr3784 | Bank of Plant Germplasm, Cuenca, Spain |
| | 160 | cr3785 | Farmer's market, Les Vans, Dept. Ardèche, France |
| *C. cartwrightianus* Herb. | | | |
| | | cr2806 | Attica, Greece |
| | | cr3275 | Attica, Greece |
| | | cr3282 | Attica, Greece |
| | | cr3294 | Attica, Greece |
| | | cr3899 | Kea, Greece |
| | | cr3915 | Kea, Greece |
| | | cr3255 | Tinos, Greece |
| | | cr3258 | Tinos, Greece |
| | | cr3269 | Crete, Greece |
| | | cr3270 | Crete, Greece |
| | | cr4458 | Naxos, Greece |
| | | cr4450 | Naxos, Greece |
| *C. oreocreticus* B.L.Burtt | | | |
| | | cr2579 | Crete, Greece |
| | | cr2675 | Crete, Greece |
